# CD95 receptor activation by ligand-induced trimerization is independent of its partial pre-ligand assembly

**DOI:** 10.1101/293530

**Authors:** C. Liesche, J. Berndt, F. Fricke, S. Aschenbrenner, M. Heilemann, R. Eils, J. Beaudouin

## Abstract

CD95 (Fas, APO-1, TNFRSF6) is a widely expressed single-pass transmembrane protein that is implicated in cell death, inflammatory response, proliferation and cell migration. CD95 ligand (CD95L, FasL, TNFSF6), is a potent apoptotic inducer in the membrane form but not when cleaved into soluble CD95L (sCD95L). Here, we aimed at understanding the relation between ligand-receptor multimerization and receptor activation by correlating the kinetics of ligand binding, receptor oligomerization, FADD (FAS-Associated via Death Domain) recruitment and caspase-8 activation inside living cells. Using single molecule localization microscopy and Förster resonance energy transfer imaging we show that the majority of CD95 receptors on the plasma membrane are monomeric at rest. This was confirmed functionally as the wild-type receptor is not blocked by a receptor mutant that cannot bind ligand. Moreover, using time-resolved fluorescence imaging approaches we demonstrated that receptor multimerization follows instantaneously ligand binding, whereas FADD recruitment is delayed. This process can explain the typical delay time seen with caspase-8 activity reporters. Finally, the low activity of sCD95L, which was caused by inefficient FADD recruitment, was not explained by the low avidity for the receptor but by a receptor clustering mechanism that was different from the one induced by the strong apoptosis inducer IZ-sCD95L. Our results reveal that receptor activation is modulated by the capacity of its ligand to trimerize it.

**Highlights:**

- At a density of less than 10 receptors per µm^2^ CD95 exists as monomer (58%) and dimer (42%)
- Pre-formed dimers do not contribute to ligand-induced CD95 apoptotic signaling
- The PLAD of CD95 attenuates overexpression-induced, ligand-independent cell death
- soluble CD95L can rapidly multimerize CD95 after binding but it is still a poor inducer of apoptosis through inefficient FADD recruitment
- FADD recruitment kinetics but not ligand binding kinetics correlates with caspase-8 onset of activity

## Introduction

Cell surface receptors translate extracellular stimuli into intracellular signals through changes in their intracellular domain that allow post-translational modifications or new protein-protein interactions. The activation of single-pass transmembrane receptors typically involves their multimerization, before and/or after ligand binding (*1*). However, our understanding of receptor activation is limited due to the challenge in getting a complete structural picture of the receptor before and after ligand binding (*2*)(*3*). Here, we aimed at better understanding the relation between multimerization and activation of the TNF receptor superfamily member CD95 (Fas, APO-1, TNFRSF6) using fluorescence-based cell biology approaches.

Upon CD95 activation, the death-inducing signaling complex (DISC) is assembled on the cytosolic side (*4*). The DISC was biochemically observed as a so-called SDS-stable aggregate of 150-250 kDa and contains the receptor, FADD, caspase-8/-10 and cFLIP isoforms (*4*)(*5*)(*6*). CD95 receptor and FADD adaptor protein each contain the so-called death domain (DD) which allows recruitment of FADD to CD95 through an homotypic interaction (*7*)(*8*). Likewise, the death effector domain (DED), which is present in FADD, procaspase-8 and −10 and cFLIP isoforms, makes homotypic interactions and thus enables recruitment of DED-containing proteins to FADD. The DED appears twice in procaspase-8 and −10 and cFLIP, and induces a chain of procaspase-8/-10, and cFLIP long isoforms (*9*)(*10*). Moreover, next to apoptosis, DISC formation can induce necrosis, NFκB or PI3K activation (*11*)(*12*) and can thus lead to different phenotypes, like cell death (*4*)(*13*)(*14*)(*15*), inflammation (*16*), cell proliferation (*17*) or migration (*18*).

CD95 ligand (CD95L, FasL, TNFSF6) is a type-II transmembrane protein principally expressed in cytotoxic T cells and natural killer cells. Those cells induce death of target cells through the granzyme/perforin pathway or through presentation of CD95L (*19*). The extracellular domain of CD95L can be cleaved by proteases (*20*)(*21*)(*22*) and the soluble product, sCD95L, is a weak inducer of apoptosis that can block the activity of the membrane form (*23*)(*20*)(*24*)(*25*). Using gel filtration (*25*), X-ray crystallography (*26*) and single molecule fluorescence microscopy (*27*) it was shown that sCD95L forms a trimer. Two types of modifications can transform sCD95L into a strong apoptosis inducer: its secondary multimerization using antibodies (*24*)(*28*) or its recombinant fusion to a trimerization domain, like the isoleucine zipper (IZ) (*29*) or the foldon that is part of the protein fibritin from the bacteriophage T4 (*30*).

Different receptor clustering mechanisms have been proposed for different stimulating agonists and cellular contexts (*11*)(*31*)(*32*). Three structural studies showed that the interactions between the DDs of CD95 and FADD may drive receptor clustering. On the one hand, a change of conformation of CD95 DD was observed and the formation of an open CD95 receptor network was proposed (*33*). On the other hand, using different protein crystallization conditions, a closed complex comprised of 5-7 receptor DD and 5 FADD DD with no change of conformation was observed (*34*)(*35*). Another potential source of receptor clustering is the modulation of its interaction with membrane lipids, as for example CD95 palmitoylation is shown to be involved in the SDS-stable aggregation (*36*)(*37*). Moreover, the signaling form of CD95 was proposed to trimerize through its transmembrane domain (*38*). Finally, the ligand-receptor interaction itself contributes to receptor clustering (*11*). The trimeric nature of CD95L and of relatives of the TNF family originally lead to the idea that it induces receptor trimerization, as it was observed by crystallization of TNFβ with the extracellular domain of TNFR1 (*39*). Later, by crystallization of the TNFR1 extracellular domain in the absence of ligand, it was shown that TNFR1 itself can form dimers and that one of the identified dimeric structures would bind ligand while staying dimeric (*40*)(*41*). Following experiments investigating the N-terminal domain of CD95 led to the identification of the so-called pre-ligand assembly domain (PLAD) that allows receptor oligomerization in the absence of ligand (*42*)(*43*)(*44*).

From that, one can propose a variety of models for the ligand-receptor interaction. If receptors are monomeric, the ligand may simply trimerize it. If they are oligomeric, then the ligand interaction may lead to the formation of a two-dimensional network (*45*)(*41*). Alternatively, one can also envision a dynamic equilibrium between monomeric and trimeric CD95, allowing the trimeric ligand to insert itself in the trimeric receptor (*11*).

To test those different interaction models, we used fluorescence microscopy approaches, allowing us in particular to characterize the stoichiometry of the receptor and the kinetics of ligand binding and receptor multimerization. Using single-molecule super-resolution microscopy to count receptors within complexes (*46*) and other functional approaches, we could observe that CD95 oligomerization in the absence of ligand is limited and does not influence its apoptotic signaling in the presence of ligand. This would be consistent with a model where receptor activation occurs through trimerization by its ligand. To get insights into CD95 signaling, we compared receptor activation by sCD95L and IZ-sCD95L. Although both ligands are trimeric at the population level, sCD95L had a lower avidity to the receptor. Moreover, even at equal receptor occupancy, sCD95L induced less FADD recruitment and less caspase-8 activity than IZ-sCD95L. Fluorescence recovery after photobleaching (FRAP) imaging, Förster resonance energy transfer (FRET) imaging and measurement of ligand unbinding kinetics showed that CD95 multimerization is performed rapidly, yet differently, by sCD95L and by IZ-sCD95L. Our results let us propose that receptor activation can be controlled by the capacity of its ligand to trimerize it.

## Results

### CD95 receptor counting on the plasma membrane using PALM shows predominantly monomeric CD95

Like TNFR, CD95 is thought to be trimerized or dimerized in the absence of ligand through an interaction driven by the PLAD, a conclusion notably motivated by biochemical crosslinking experiments and FRET data (*43*)(*44*). We previously showed that we can identify the stoichiometry of receptors on the plasma membrane with high precision using photoactivated localization microscopy (PALM) (*46*). By counting blinking events from mEos2 fusions and analyzing their statistics with kinetic models (*47*), we could identify the oligomeric state of the trimeric Vesicular stomatitis New Jersey virus Glycoprotein G (VSV-G), the dimeric Cytotoxic T-lymphocyte protein 4 (CTLA-4) and the monomeric T-lymphocyte activation antigen CD86 (*46*). Here we used this approach to quantify the oligomerization state of CD95 on the plasma membrane of HeLa cells. We used CRISPR/Cas9 to generate a monoclonal population of HeLa(CD95^KO^) cells, which lack expression of endogenous CD95 (**Fig. 1A**) and for which we could restore apoptosis sensitivity by expressing exogenous CD95-mGFP (**Fig. S1**). We next expressed mEos2-labeled CD95 in HeLa(CD95^KO^) cells. At a density of less than 10 receptors per µm^2^, we observed a punctate pattern on the cell surface (**Fig. 1B**). We performed single molecule localization microscopy and analyzed the distribution of the number of blinking events in each localization cluster as in (*47*). Assuming a monomer-dimer mixture, this model allowed determining a fraction of 58% CD95 monomers and 42% CD95 dimers (**Fig. 1B, histogram**). These data also showed that CD95 at rest does not exclusively pre-oligomerize.

**Fig. 1.**
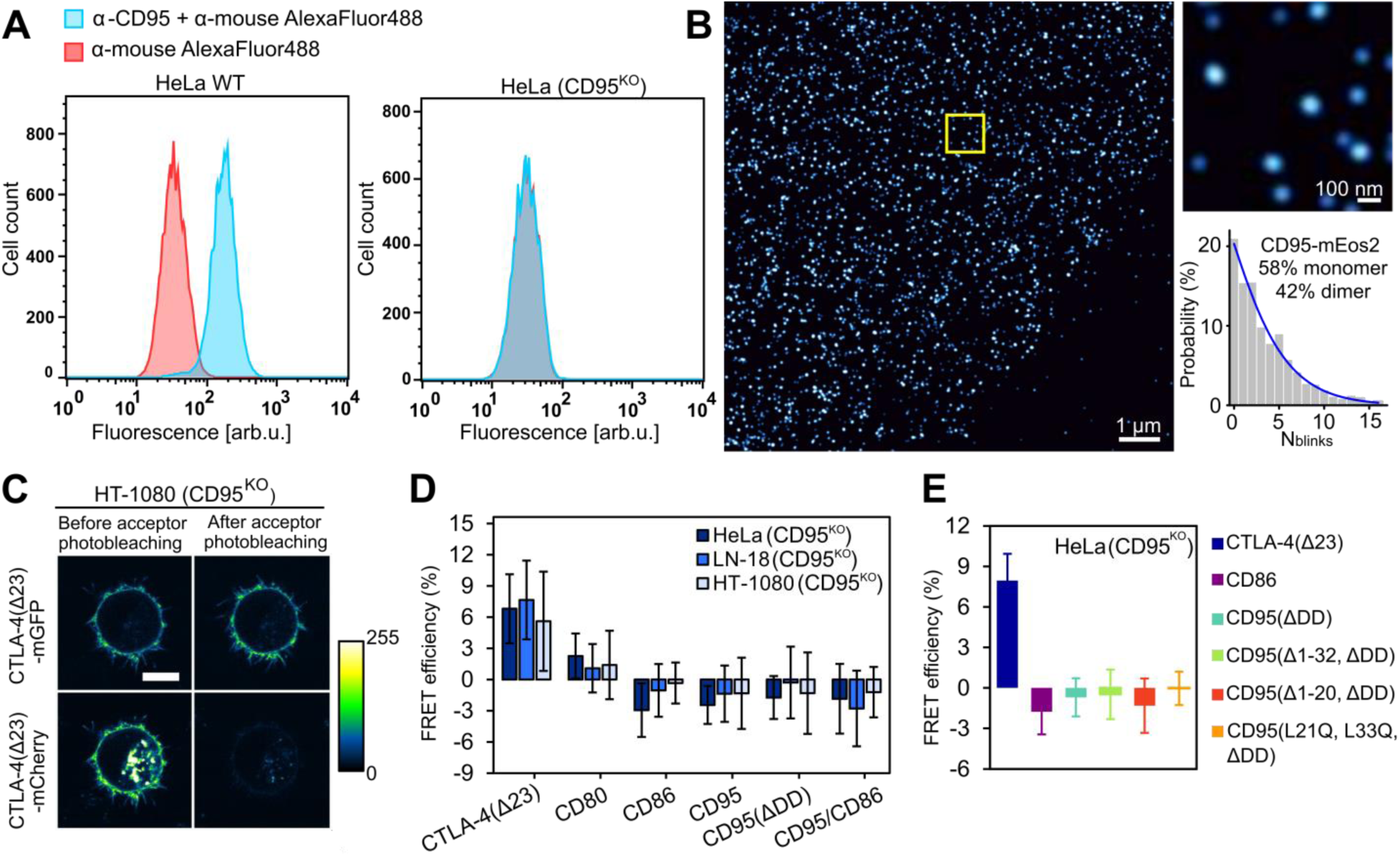
The majority of CD95 receptor on the plasma membrane is monomeric. (**A**) HeLa wild-type and HeLa(CD95^KO^) cells were stained by immunofluorescence using APO-1-3 antibody and an Alexa488 secondary antibody, or using only the secondary antibody as negative control. The overlapping histograms for HeLa(CD95^KO^) cells show the absence of CD95 protein. (**B**) PALM image of CD95-mEos2 at the membrane of HeLa(CD95^KO^) cells. The distribution of blink number per localization cluster revealed a fraction of 58% monomers and 42% dimers (n= 14 cells). (**C**) Confocal microscopy images show CTLA-4(Δ23)-mCherry and CTLA-4(Δ23)-mGFP at the plasma membrane before and after acceptor photobleaching in HT-1080 (CD95^KO^) cells. Scale bar 10 µm. (**D**) FRET acceptor photobleaching data in HeLa(CD95^KO^) (dark blue), LN-18(CD95^KO^) (blue) and HT-1080(CD95^KO^) cells (light blue). Pairs of mGFP and mCherry-tagged proteins (CTLA-4(Δ23), CD80, CD86, CD95 or CD95(ΔDD)) were co-expressed from a single open reading frame using the 2A peptide. The monomeric receptor CD86 and a pair of CD86-mCherry and CD95-mGFP served as negative controls. (**E**) Pairs of mGFP-and mCherry-tagged proteins (CTLA-4(Δ23), CD86, CD95(ΔDD), CD95(Δ1-32, ΔDD), CD95(Δ1-20, ΔDD) or CD95(L21Q, L33Q, ΔDD)) were co-expressed in HeLa(CD95^KO^) cells using the 2A peptide. Absence of FRET is observed for CD95 receptors at rest.

Our results may oppose previous studies (*43*)(*44*) that led to the model of oligomeric CD95 in the absence of ligand. Therefore, we revisited two of the original experiments that motivated the oligomerization model. First, FRET measurements of overexpressed receptor fusions and second, the influence on cell death of a receptor mutant, CD95(R86S), that has an intact PLAD but that lacks ligand interaction sites.

### Unliganded CD95 receptors on the plasma membrane do not show FRET

To test CD95 receptor oligomerization by FRET imaging, we set-up acceptor photobleaching experiments in three different tumor cell lines: HeLa, the glioblastoma cell line LN-18 and the fibrosarcoma cell line HT-1080. We used CRISPR/cas9 to generate clones that lack CD95 (**Fig. S2A**). To stay consistent with the other experiments (see below), we chose mGFP and mCherry as FRET pair and fused them to the cytosolic C-terminus of CD95, termed CD95-mGFP and CD95-mCherry. In order to get equal expression of both fusions, we co-expressed them from a single open reading frame carrying the coding sequence of the 2A peptide of *Thosea Asigna* virus in between (*48*). The control constructs mGFP-2A-mCherry and mCherry-2A-mGFP showed that the amount of protein encoded after the 2A peptide was 0.8 times the one of the protein encoded in front (**Fig. S2B**). As positive control for FRET, we used the dimeric plasma membrane protein CTLA-4(Δ23), a mutant of CTLA-4 that lacks the last 23 amino acids to favor its localization at the plasma membrane (*49*). A second positive control for FRET was the plasma membrane receptor CD80, which was shown to partially dimerize (*50*)(*46*)(*47*). The monomeric receptor CD86 and a pair of CD86-mCherry and CD95-mGFP served as negative control. Next to wild-type CD95 receptor, we also measured a truncation mutant of CD95, termed CD95(ΔDD), comprising the first 210 amino acids but lacking the death domain. To note, cells with similar fluorescence intensities were selected for FRET measurements (**Fig. S3A**). In particular, wild-type CD95 receptors did not induce spontaneous cell death at those intensities (see below). Moreover, contrary to HeLa(CD95^KO^), adherent LN-18(CD95^KO^) and HT-1080(CD95^KO^) cells moved during the measurement, making the comparison of mGFP signal before and after photobleaching unreliable. This problem was solved by imaging trypsinized cells and focusing in the middle of the cell (**Fig. 1C**). The measurement of a batch of cells was limited to 20 minutes to minimize trypsin exposure. FRET was measured for each cell after segmentation of the plasma membrane signal (please see **supplemental info**). For each tested receptor construct, we obtained comparable FRET signals among the three cells lines (**Fig. 1D**): Highest FRET signals were measured for CTLA-4(Δ23) receptors followed by CD80. CD86, CD95 and CD95(ΔDD) showed no FRET in the three tested cell lines. Signals with CD95 or CD95(ΔDD) were not significantly different from those obtained with CD86 or CD95 combined with CD86 but they were significantly different from the one obtained with CTLA-4(Δ23) (**Fig. 1D, and Supplementary Table 1**).

As the absence of FRET appeared in contradiction with previous studies (*43*)(*44*), we tested the possibility that the PLAD was cleaved in those cell lines. The idea was motivated by the observation in (*51*) that the matrix metalloproteinase-7 (MMP-7) can cleave the N-terminal part of CD95 in front of leucine at position 20 and 32, with the numbering starting at amino acid 18 in the signal peptide of CD95 (**Fig. S4A**). In line, CD95-mGFP and CD95(ΔDD)-mGFP appeared as a doublet by western blot, potentially reflecting this cleavage (**Fig. S4B**). We designed mGFP and mCherry-fused CD95(L21Q,L33Q) and CD95(L21Q,L33Q, ΔDD) receptors in which leucine 21 and 33 are mutated, and receptor truncations starting at leucine 21 and leucine 33 respectively CD95(Δ1-20) and CD95(Δ1-32). While CD95(Δ1-20) and CD95(Δ1-32) truncations resembled the size of the corresponding cleavages, the higher band of the doublet predominated for the double mutant CD95(L21Q,L33Q) in CD95 or CD95(ΔDD) transfected cells (**Fig. S4C**), confirming the involvement of Leu20 and Leu32 in CD95 cleavage as described in (*51*). Still, FRET was neither observed with fluorescent protein fusions of CD95(L21Q,L33Q) nor with CD95(Δ1-20) or CD95(Δ1-32), suggesting that the absence of FRET for CD95 receptors was not due to truncated PLAD (**Fig. 1E**). These data support our results obtained from PALM measurements in that the majority of CD95 is monomeric.

### The receptor mutant CD95(R86S) does not function as dominant negative receptor

The second indication for CD95 oligomerization in the absence of ligand was that two receptor mutants would block the activity of the wild-type receptor in a dominant negative manner (*44*). These mutants were CD95(R86S), a receptor that fails to bind CD95L, and CD95(ALPS Pt2), a receptor that lacks amino acids 52 to 96 corresponding to exon 3. In order to reproduce and further quantify potential dominant negative effects, we co-expressed receptors using the 2A peptide and measured the impact of this co-expression on ligand-induced cell death. For this, we placed the mCherry-fused receptor CD95(R86S), CD95(ALPS Pt2), CD95(ΔDD) or CD86 in front of the 2A peptide and mGFP-fused wild-type CD95 receptor after it (**Fig. S5A-E**). Of note, all tested receptors displayed fluorescence at the plasma membrane except CD95(ALPS Pt2)-mCherry, which was absent at the plasma membrane but present inside the cell with a pattern reminiscent of the endoplasmic reticulum (**Fig. S5C**). Therefore, this receptor unlikely blocks wild-type receptor activity at the plasma membrane. Next, using confocal microscopy, we confirmed that CD95(R86S)-mGFP, but not CD95-mGFP, fails to bind mCherry-IZ-sCD95L (**Fig. S5F**). The cell lines HeLa(CD95^KO^), LN-18(CD95^KO^) and HT-1080(CD95^KO^) were each transfected with the different receptor mutants. Cells were stimulated 72 h later with IZ-sCD95L and imaged by confocal time-lapse microscopy. We analyzed the time of death of transfected cells, recognized from the mGFP signal, by identifying the time point where cell rounding characteristic for cell death occurred. Compared to the control situation with CD86-mCherry, co-expression of CD95(ΔDD)-mCherry with CD95-mGFP lead to a drastic decrease of cell death showing a block of CD95-mGFP activity by CD95(ΔDD)-mCherry (**Fig. 2A**). However, no difference in the time of death was observed when CD95-mGFP was co-expressed with CD95(R86S)-mCherry or CD95(ALPS Pt2)-mCherry, unambiguously showing that neither of the two proteins blocks CD95-mGFP activity. To further validate our results, we designed a similar experiment with unlabeled receptors. To recognize transfected cells we placed an internal ribosome entry site (IRES) and the GFP-encoding gene on the same plasmid (see scheme **Fig. 2B**). These constructs allowed a correlation between the time of death upon IZ-sCD95L incubation and the GFP fluorescence signal. As expected, cells with higher GFP signals, and therefore with higher receptor levels, died earlier than cells with lower GFP fluorescence signal (**Fig. 2B**). Furthermore, consistent with the fluorescent protein fusions shown in **Fig. 2A**, co-expression of CD95(ΔDD) with CD95 led to an increase of the time of death, while no difference in the correlation of the time of death and the GFP signal was observed with the receptor pairs CD95/CD86, CD95/CD95(R86S) and CD95/CD95(ALPS Pt2). Taken together, our data clearly show that CD95(R86S) and CD95(ALPS Pt2) do not act as dominant negative mutants.

**Fig. 2.**
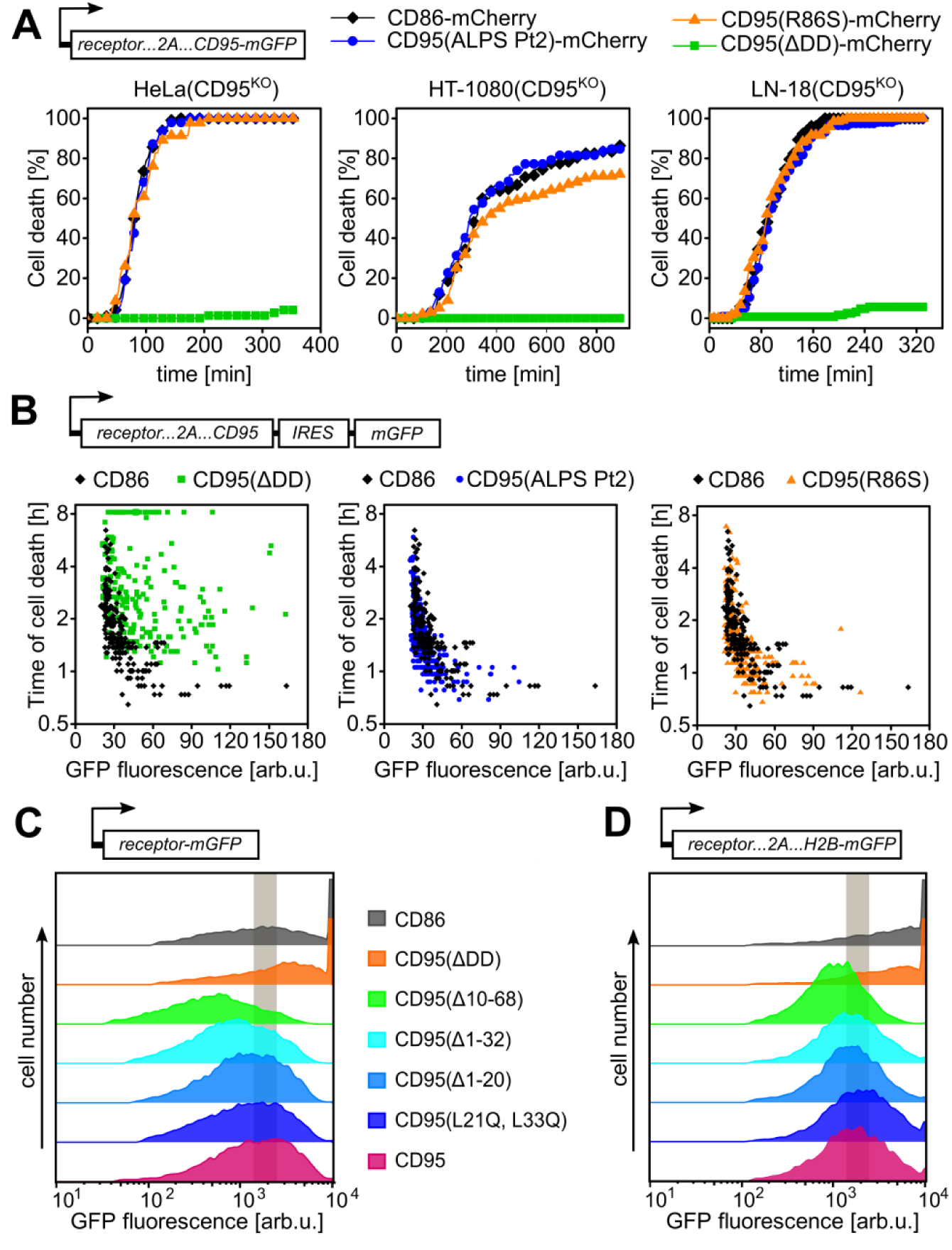
CD95(R86S) does not function as a dominant negative receptor, but the N-terminal domain of CD95 reduces CD95 receptor self-activation. (**A**) mCherry-fused CD95(R86S), CD95(ALPS Pt2), CD95(ΔDD) and CD86 were coexpressed with mGFP-fused wild-type CD95 in HeLa(CD95^KO^), HT-1080(CD95^KO^) and LN-18(CD95^KO^) for 72 h. Cells were incubated with 100 ng/ml (for HeLa) or 1 ng/ml (for LN-18 and HT-1080) IZ-sCD95L and monitored by confocal microscopy, n = 47 to 98 fluorescent cells were analyzed per condition. (**B**) CD95(R86S), CD95(ALPS Pt2), CD95(ΔDD) or CD86 were coexpressed with wild-type CD95 and mGFP in HeLa(CD95^KO^) cells for 72 h. The time of cell death (log2) is plotted against the mGFP intensity measured before induction. Each condition shows n = 202 to 319 cells. (**C**) 200 ng plasmid DNA encoding mGFP fusions of CD86, CD95(ΔDD), and the different N-terminal CD95 mutants and truncations were transfected in 5×10^4^ cells. Histograms are flow cytometry data showing mGFP fluorescence in propidium iodide-positive cells. Highest mGFP fluorescence amounts were observed with the controls CD86 and CD95(ΔDD) and the lowest with CD95(Δ1-68). (**D**) Same workflow as in C, using unlabeled receptors and coexpression with H2B-mGFP as expression read-out.

### The CD95 N-terminal domain controls receptor self-activation

CD95 self-interaction was originally studied by FRET using CD95 fused to fluorescent proteins in HEK 293T cells (*44*). However, this FRET signal did not linearly increase with receptor expression like a positive control but appeared only in cells which expressed high amounts of receptor (*44*). Of note, HEK 293T cells do not die from CD95 apoptosis and thus allow very high CD95 expression. In view of our results described above, we hypothesized that the high expression needed to observe FRET in HEK 293T cells may induce spontaneous cell death through CD95 self-interaction in apoptosis-sensitive cells. To test this, we measured the amount of expressed mGFP-tagged receptors in dead cells in the absence of ligand. For this assay, we transfected HeLa(CD95^KO^) with up to four times more DNA than in experiments shown in **Fig. 2**, stained cells 72 h later with propidium iodide (PI) and measured fluorescence by flow cytometry. Of note, increasing plasmid concentration led to a saturation of the mGFP signal in PI-positive cells, showing that the receptor amount was not underestimated by limited amount of DNA (**Fig. S6).** While CD95(ΔDD)-mGFP and the control construct CD86-mGFP reached similar GFP fluorescence in PI-positive cells, CD95-mGFP transfected cells displayed 10 times less GFP fluorescence (**Fig. 2C**). Likewise, PI-positive cells expressing the receptor mutants related to MMP7, CD95(L21Q,L33Q)-mGFP, CD95(Δ1-20)-mGFP and CD95(Δ1-32)-mGFP, showed the same GFP fluorescence intensities as wild-type CD95. This clearly shows that CD95 self-activates when massively over-expressed to induce death in a ligand-independent manner. The PLAD domain of CD95, which has self-interacting properties, was mapped in the CRD1 of the receptor (*52*). To test its impact on CD95 self-activation, we designed the truncation mutant CD95(Δ10-68)-mGFP that lacks the entire N-terminal part of CD95 including the CRD1, from amino acids 10 to 68. We found that this mutant was expressed four-fold less compared to the wild type CD95-mGFP when cell death occurred, showing that lack of this N-terminal part of the receptor facilitates spontaneous CD95 self-activation at this high receptor expression. In other words, the PLAD protects cells from this type of ligand-independent cell death (**Fig. 2C**). Equivalent observations were made when we performed the experiment with unlabeled receptors and using H2B-mGFP as expression readout (**Fig. 2D**). In conclusion, we propose that CD95 self-activation, which can occur at increased receptor amounts in apoptosis-sensitive cells, is attenuated by the N-terminal domain of CD95.

Taken together, using counting of receptors by PALM, FRET imaging and death kinetic analysis of co-expressed CD95 mutants, we showed that the majority of CD95 does not exclusively pre-associate but that the monomeric form of CD95 predominates on the plasma membrane of cells at rest.

### Equal CD95 receptor occupancy with sCD95L and IZ-sCD95L translates into different caspase-8 activity

Since our data strongly indicated that CD95 is monomeric at rest and that the pre-oligomerization is not influencing apoptosis signaling, the receptor is likely activated through trimerization by CD95L. Such a process implies a multi-step complex formation, starting with binding of the ligand to a first receptor on the plasma membrane, and followed by the recruitment of a second and a third receptor to form a 1:3 trimeric ligand:receptor complex. To get insights into this process, we compared two different recombinant ligands, the natural soluble CD95L (sCD95L), a weak apoptosis inducer (*23*), and the isoleucine zipper-fused ligand (IZ-sCD95L), which is a strong apoptosis inducer (*29*)(*53*). We have previously shown that sCD95L and IZ-sCD95L are trimeric at the population level even in the picomolar range (*27*), a result that we validated here by comparing the brightness of mCherry-sCD95L, mCherry-IZ-sCD95L and mCherry-IZ to the one of mCherry alone by fluorescence correlation spectroscopy (FCS) (**Fig. S7A**). mCherry-tagged sCD95L and IZ-sCD95L were expressed through secretion by HEK293T cells at concentrations between 1 and 5 µg/ml, as assessed by FCS (**Fig. S7B-D**). To quantify ligand binding, we used mGFP-labeled CD95 receptors stably expressed in HeLa cells (HeLa(CD95-mGFP)). Receptor occupancy was quantified by segmenting the plasma membrane signal (**see supplemental info for details**) and by calibrating the mCherry signal to the mGFP one (**Fig. S7E**). As the mCherry signal was also visible in the medium next to the cells, cells were briefly washed before measurement to get a precise measurement of the receptor occupancy at a given time point (**Fig. S7F**). By quantifying ligand binding after 10 min using different ligand doses, we found that about 3.5 fold more mCherry-sCD95L than mCherry-IZ-sCD95L was required to reach the same receptor occupancy (**Fig. 3A**). To test if this difference in affinity was enough to explain their different activity, we next measured cell death in HeLa(CD95-mGFP) cells incubated with 1.8 µg/ml mCherry-IZ-sCD95L or with 6.4 µg/ml mCherry-sCD95L. While both ligands reached equal receptor occupancies with similar kinetics (**Fig. S7G**), 56 min were required to reach 50% cell death with mCherry-IZ-sCD95L while 2h 23 min were required with mCherry-sCD95L (**Fig. 3B**). In line, caspase-8 activity, measured with a localization reporter (**Fig. 3C**) as previously described in (*54*), showed a weaker cleavage rate and a larger onset with mCherry-sCD95L than with mCherry-IZ-sCD95L (**Fig. 3D and Fig. S8**). These data show that equal CD95 receptor occupancy with sCD95L and IZ-sCD95L does not translate into equal caspase-8 activity. sCD95L and IZ-sCD95L activity was further explored over a wider range of ligand concentrations and for two different receptor levels. Target cells were wild-type HeLa and HeLa stably overexpressing CD95 (HeLa(CD95)) that have about 12000 and 160000 receptors, respectively (*55*). We here used labeled and unlabeled ligand forms and we estimated the concentration of the unlabeled ligand by comparison of cell death to the labeled ligand (**Fig. S7H**). Contrary to IZ-sCD95L, sCD95L poorly induced apoptosis in HeLa cells (**Fig. 3E**). However, sCD95L efficiently induced apoptosis in HeLa(CD95) cells. Moreover, increasing ligand concentrations led to a decrease in the median cell death time, which was saturating at 60 min with IZ-sCD95L in HeLa cells and at 38 min with HeLa(CD95) cells. The minimal time required by sCD95L to kill HeLa(CD95) was around 70 min, which means that increasing sCD95L concentration could not compensate for its weak activity, even at high receptor amounts.

**Fig. 3.**
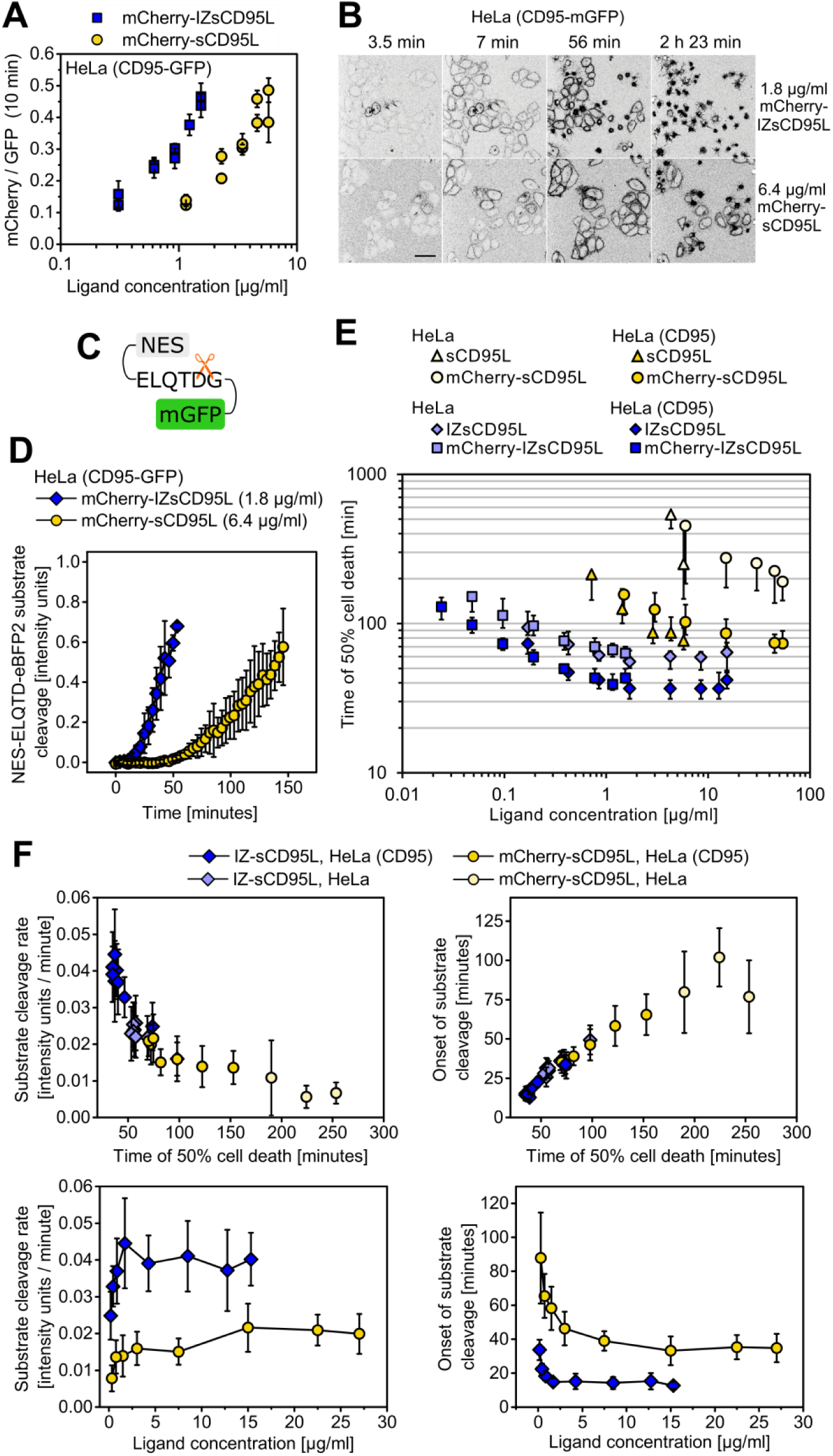
Equal receptor occupancy with sCD95L and IZ-sCD95L does not translate into equal caspase-8 activity. (**A**) HeLa(CD95-mGFP) cells were incubated with mCherry-fused ligands and washed after 10 min to quantify the receptor occupancy (ratio of mCherry to mGFP signal). (**B**) Fluorescence microscopy images showing mCherry-IZ-sCD95L (1.8 µg/ml) and mCherry-sCD95L (6.4 µg/ml) binding to HeLa(CD95-mGFP) cells. Note the different time of death. Scale bar 50 µm. (**C**) Caspase-8 activity reporter, NES: nuclear export signal. Cleavage of the peptide linker ELQTDG by caspase-8 redistributes mGFP over the cell. (**D**) Caspase-8 activity measurement using the reporter NES-ELQTDG-eBFP2 in HeLa(CD95-mGFP) cells stimulated as in (B). (**E**) Median cell death times are plotted with 25th and 75th percentiles as a function of ligand concentration. Each data point corresponds to 150 to 250 cells. (**F**) Upper row: rate and onset of substrate cleavage as a function of time of death. Note the correlation independent of ligand form and receptor amount. Lower row: rate and onset of substrate cleavage in HeLa(CD95) as a function of ligand concentration. Note that the saturation reached by both ligands is at different rates and onsets.

To test how these results were related to DISC activity, we also measured caspase-8 substrate cleavage (**Fig. 3F**). For this, we extracted the rate and onset from caspase-8 activity data by fitting progress curves from single cell measurements to the modified Gompertz function (*56*) (**Fig. S8**). The correlation of substrate cleavage rate or onset of activity with the time of cell death coincided for the two different ligands and the two different receptor amounts showing that mechanisms downstream of caspase-8 activation were the same for both ligands (**Fig. 3F**). Of note, at saturating ligand concentrations, caspase-8 substrate cleavage rates in HeLa(CD95) cells were two times smaller with mCherry-sCD95L than with IZ-sCD95L showing that mCherry-sCD95L was not able to trigger full caspase-8 activity. Likewise, the onset of caspase-8 activity was about two times later for sCD95L (35 min) than for IZ-sCD95L (15 min) demonstrating that even in saturating conditions caspase-8 activation by sCD95L is limited by a process upstream of caspase-8.

### Rapid ligand binding is followed by slow FADD recruitment

FADD is the bridging protein between activated receptors and caspase-8 in the DISC. To assess its recruitment kinetics to the plasma membrane, we generated a HeLa(CD95) cell line that lacks endogenous FADD and that stably expresses FADD-mGFP (**Fig. S9**). Monoclonal cell lines were selected, each showing a relatively large cell-to-cell variability in FADD expression. We then stimulated clones with IZ-sCD95L and imaged mGFP over time using confocal microscopy. We found cells with clear FADD-mGFP signal at the plasma membrane increasing over time (**Fig. 4A and movie 1**). As we could not easily analyze the plasma membrane signal, we quantified FADD recruitment by measuring the decrease in mean FADD-mGFP intensity inside the cell over time. The relative decrease in fluorescence was determined by normalizing the data to the fluorescence intensity before addition of ligand. The kinetics of FADD-mGFP recruitment upon stimulation with IZ-sCD95L was reproducible in each of the tested cell lines (**Fig. 4B**). Remarkably, it displayed a lag phase of 10 min showing that FADD recruitment does not immediately follow receptor binding. To compare FADD-mGFP recruitment with sCD95L and IZ-sCD95L, we induced two different FADD-mGFP expressing cell clones with mCherry-sCD95L (2.5 µg/ml) and mCherry-IZ-sCD95L (0.9 µg/ml) in presence of 50 µM zVAD to prevent cell death. At these concentrations of ligand, receptors showed equal occupancy with ligand (**Fig. 4C**). It should be noted that upon stimulation we observed formation of large FADD-mGFP aggregates inside the cell at late time points (**Movie 2**). Those aggregates appeared in cells with a strong initial FADD-mGFP signal and we excluded them from the analysis in order to not bias the results. For mCherry-IZ-sCD95L, the lag phase of about 10 min was followed by a 20% decrease in fluorescence within 30 min and a 30% decrease within 60 min (**Fig. 4D**). In contrast, FADD-mGFP recruitment in mCherry-sCD95L induced cells was weak with only 7% within 60 min. Thus, sCD95L is not able to induce efficient FADD recruitment despite strong receptor binding.

**Fig. 4.**
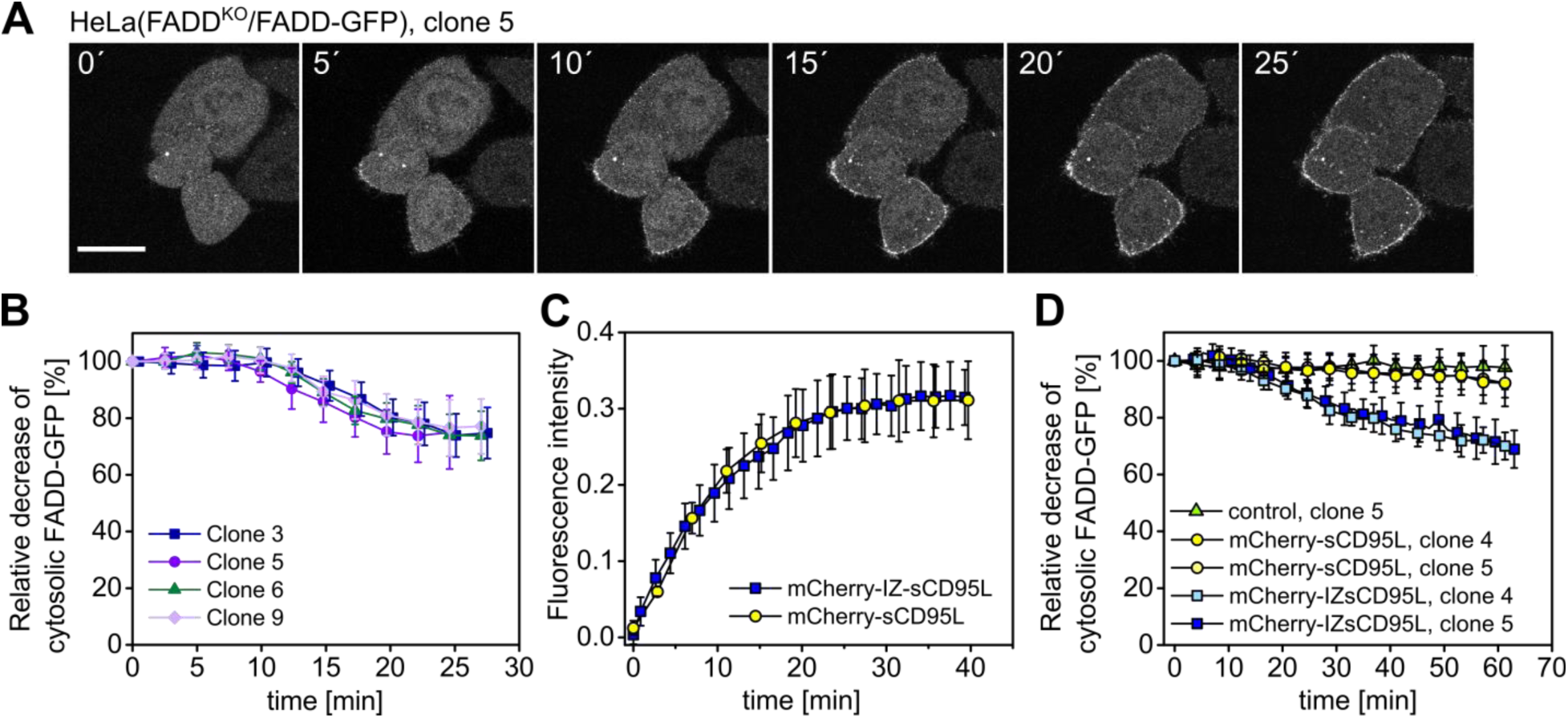
FADD recruitment to the plasma membrane shows a lag phase, whereas ligand binding occurs instantaneously. (**A**) HeLa(FADD^KO^) cells stably expressing FADD-mGFP were stimulated with IZ-sCD95L. FADD-mGFP fluorescence at the plasma membrane increases over time while the intracellular signal decreases. Images are before cell death. Scale bar 20 µm. (**B**) Quantification of intracellular FADD-mGFP over time for four different stable cell lines stimulated with IZ-sCD95L as in (**A**). Mean and standard deviation of 10, 18, 23 and 25 cells. (**C**) Binding of 2.5 µg/ml mCherry-sCD95L and 0.9 µg/ml mCherry-IZ-sCD95L to HeLa(FADD^KO^)/FADD-mGFP (clone 4) in presence of 50 µM z-VAD. mCherry signal was quantified from the whole image, subtracted with signal between cells and normalized to the number of cells on the first frame. The rate of fluorescence signal increase is maximal at time zero. Mean and standard deviation of 5 fields of view with 8 to 54 cells per field. (**D**) Intracellular FADD-mGFP fluorescence decrease of HeLa(FADD^KO^)/FADD-mGFP cells (clone 4 or clone 5) upon incubation with 2.5 µg/ml mCherry-sCD95L and 0.9 µg/ml mCherry-IZ-sCD95L in presence of 50 µM z-VAD. Mean and standard deviation of 12 to 25 cells per condition.

### The different apoptotic activity of sCD95L and IZ-sCD95L correlates with different receptor multimerization

We showed that sCD95L weak apoptotic activity is not explained by a weak avidity for plasma membrane receptors and we therefore hypothesized that receptor multimerization by sCD95L does not favor receptor activation. To test this, we first measured unbinding kinetics of mCherry-labeled ligands in the absence and in the presence of unlabeled ligand. In a ligand-dependent receptor trimerization model, the labeled ligand, visible as mCherry signal on the plasma membrane, appears as unbound only when it dissociates from all (up to three) bound receptors. When ligand unbinds one receptor but stays bound to one or two other receptors, it can very likely re-bind another free receptor. In contrast, in presence of excess unlabeled ligand, rebinding of free receptors by the labeled ligand is unlikely because unlabeled ligand competes for free receptors on the plasma membrane. This behavior could be observed with both ligands: when the supernatant of cells containing mCherry-tagged ligand was replaced by medium, more than 80% of mCherry-signal remained on the plasma membrane after 2 hours (**Fig. 5A**). In contrast, when mCherry-tagged ligand was replaced by the respective unlabeled ligand at concentrations leading to same cell death (**Fig. 5B**), only 50% of mCherry-IZ-sCD95L and 25% of mCherry-sCD95L remained after 2 hours on cells (**Fig. 5A, upper row**). It is noteworthy that at these time scales we could clearly observe ligand internalization (**Fig. S10A**). Thus, signal loss at the plasma membrane can be due to ligand unbinding from receptors or due to ligand internalization. To discriminate between these two processes, we blocked internalization using 3 mM methyl-β-cyclodextrin (mβcd) (**Fig. S10A**). Ligand internalization has been proposed to enhance or slow down apoptosis depending on the cell type (*57*)(*58*)(*59*). Here, mβcd treatment reduced ligand internalization but also amplified caspase-8 activity and accelerated cell death (**Fig. S10B**). Still, with mβcd, mCherry-IZ-sCD95L and mCherry-sCD95L showed higher residence times in the absence of the competing unlabeled ligands than in their presence (**Fig. 5A, lower row**). Moreover, unbinding of mCherry-IZ-sCD95L was slower than of mCherry-sCD95L in presence of the unlabeled counterpart (**Fig. 5B**). This suggests a tighter receptor multimerization with IZ-sCD95L than with sCD95L.

**Fig. 5.**
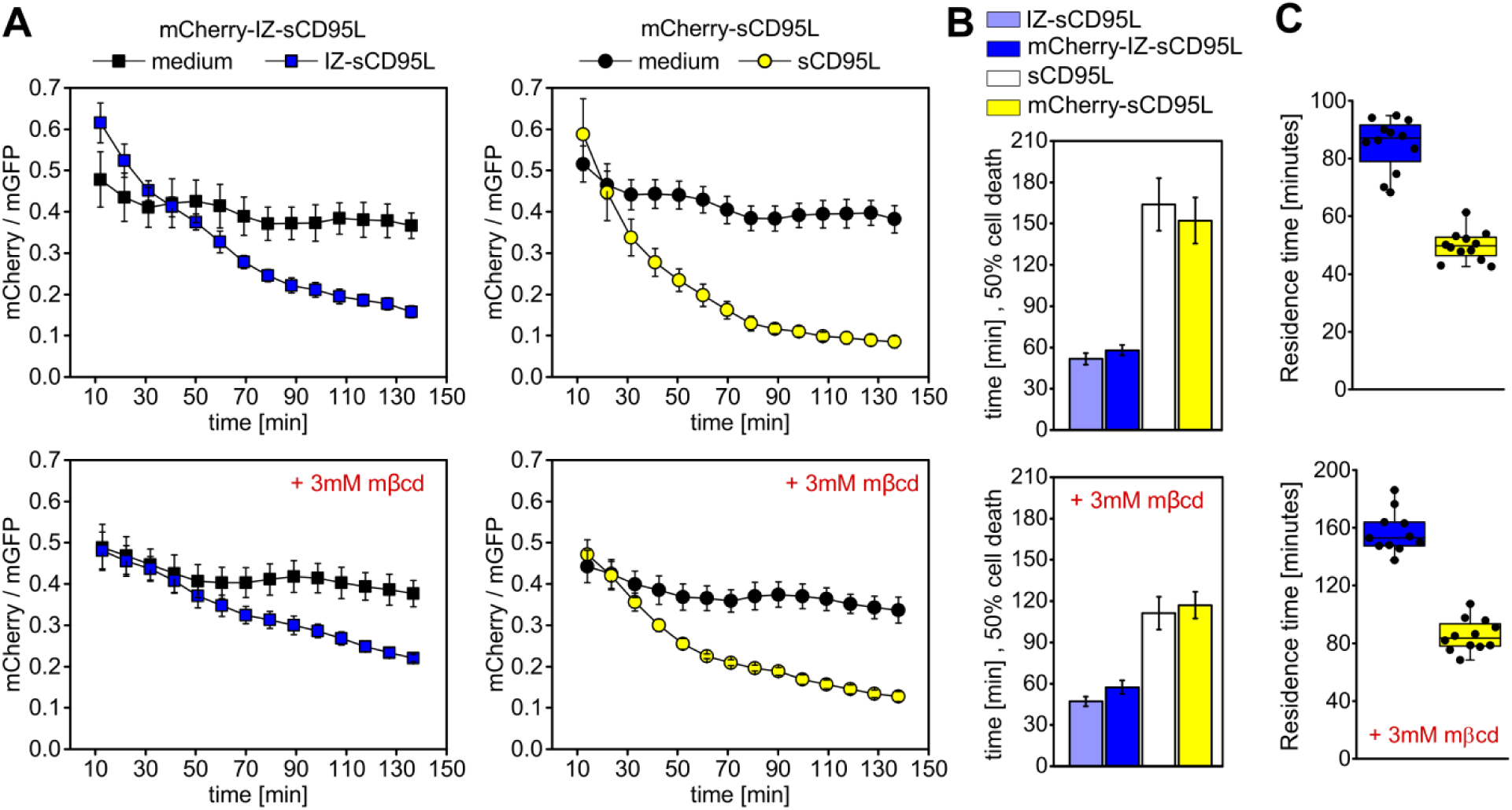
The unbinding of mCherry-sCD95L and mCherry-IZ-sCD95L from the plasma membrane is faster in presence of unlabeled competitor ligand. (**A**) Unbinding kinetics of mCherry-sCD95L and mCherry-IZ-sCD95L after washing (black circles or squares) or washing followed by addition of unlabeled ligand (yellow or blue), without (upper row) or with (lower row) mβcd. Mean and standard deviation of 12 fields of view per condition. (**B**) Time of death of HeLa(CD95) cells with (upper row) or without 3 mM mβcd and with the ligand concentrations used in (**A**). Note that in both cases, each mCherry labeled ligand and its unlabeled correspondent killed cells at the same time. (**C**) Characteristic time of unbinding of mCherry-labeled ligand following washing combined with addition of unlabeled ligand. The unbinding of ligand was faster with sCD95L than with IZ-sCD95L in absence (upper row) and in presence (lower row) of mβcd.

The second approach aimed at estimating receptor oligomerization and consisted in quantifying the diffusion coefficient of receptors upon ligand binding. We assumed that oligomerization would slow down diffusion through the increase of the number of transmembrane domains diffusing together. To measure this, we performed FRAP experiments by bleaching a thin stripe on the lower part of the plasma membrane of cells and fitting the diffusion coefficient from the broadening of the stripe over time (**Fig. 6A-B** and **see supplemental information)**. Experiments were first performed on HeLa(CD95-mGFP) cells. The estimated diffusion coefficient of non-induced receptors remained constant over time between 0.25 µm^2^/s and 0.3 µm^2^/s (**Fig. 6C**). mCherry-sCD95L and mCherry-IZ-sCD95L concentrations were at 1.8 µg/ml and 6.4 µg/ml, respectively, so that the amount of ligand bound to the cells was the same. Upon addition of ligand, the diffusion coefficient dropped by a factor of two, over 45 minutes with mCherry-IZ-sCD95L and over 110 minutes with mCherry-sCD95L. To test whether the decrease in the receptor diffusion coefficient is due to crosslinking by ligand or due to DISC assembly, we also performed the experiment using CD95(ΔDD)-mGFP transiently expressed in HeLa cells (**Fig. 6D**). To compare to CD95-mGFP, we selected cells with similar level of expression, knowing that the endogenous receptor represented about 10% of the total amount of receptors. The diffusion coefficient of CD95(ΔDD)-mGFP declined upon mCherry-IZ-sCD95L and mCherry-sCD95L to 0.09 µm^2^/s and 0.13 µm^2^/s respectively, demonstrating that the DISC formation does not contribute to the diffusion coefficient decrease. Although these FRAP experiments could indicate that sCD95L and IZ-sCD95L have a different capacity to crosslink receptors, this observation should be taken with caution because the cell-to-cell variability was high.

**Fig. 6.**
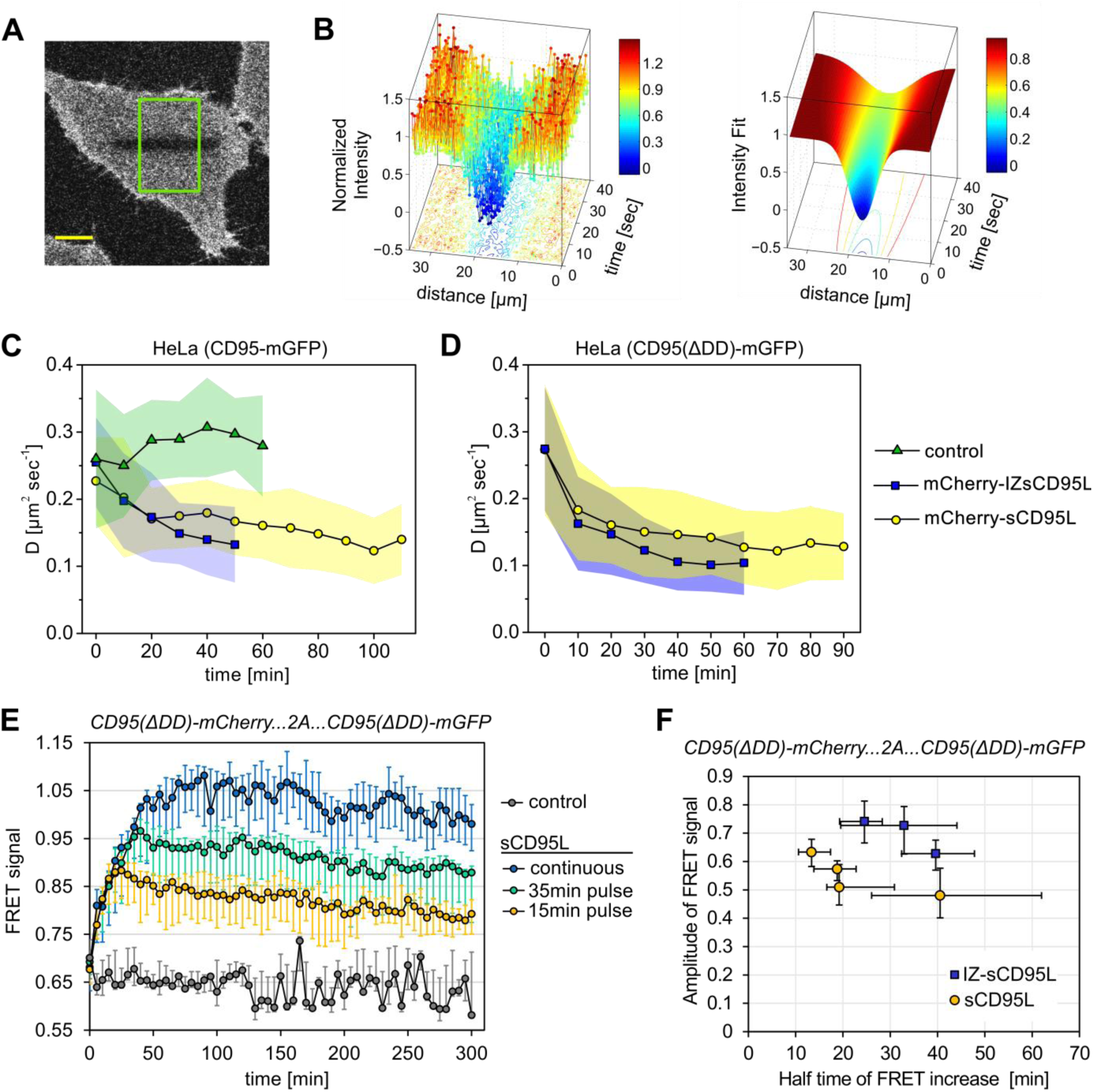
CD95 receptor is differentially crosslinked by soluble CD95L and IZ-sCD95L. (**A**) Photobleaching stripe on a HeLa(CD95-mGFP) cell. The green rectangle is a typical selection for FRAP analysis. Scale bar 10 µm. (**B**) Plot of image intensity data (left) and fit (right) from a stripe-FRAP measurement. (**C**) Diffusion coefficient (mean and standard deviation) of CD95-mGFP upon stimulation of HeLa(CD95-mGFP) with 6.4 µg/ml mCherry-sCD95L (54 cells) or 1.8 µg/ml mCherry-IZ-sCD95L (39 cells) or when left untreated (control, 25 cells). (**D**) Same with HeLa cells expressing CD95(ΔDD)-mGFP (mCherry-sCD95L, 26 cell; mCherry-IZ-sCD95L, 25 cells). (**E**) FRET (median, 25th and 75th percentiles) between CD95(ΔDD)-mGFP and CD95(ΔDD)-mCherry upon different exposure with sCD95L (continuous, 14 cells; 15 min, 15 cells; 35 min, 17 cells) or without stimulation (control, 10 cells). (**F**) FRET kinetics obtained with 1.1, 0.90, 0.67 or 0.42 µg/ml sCD95L and 0.92, 0.73 or 0.48 µg/ml IZ-sCD95L. The relation of the amplitude of the FRET signal and the half time of the FRET increase shows no overlap between the two ligands.

The third approach consisted in measuring FRET by sensitized emission between tagged receptors over time upon ligand addition using the construct CD95-ΔDD-mCherry-2A-CD95-ΔDD-mGFP expressed in HeLa(CD95^KO^). Both ligands induced a clear increase of FRET, in particular with a maximum slope at time zero similar to ligand binding kinetics (**Fig. 6E** and **Fig. S11**). Furthermore, removal of ligand after 15 min or 35 min immediately stopped FRET increase (**Fig. 6E**). Together, this indicates that receptor trimerization occurs quickly after ligand binding. To compare the multimerization capacity of both ligands, we applied different dilutions of unlabeled IZ-sCD95L and sCD95L in the range of 1 µg/ml. By parameterizing the kinetics of the FRET increase for each single cell (**Fig. S11 and supplemental material and methods**), we found that the correlation between the amplitude and the half time of FRET increase did not overlap for IZ-sCD95L and sCD95L (**Fig. 6F**). For a similar kinetics, the amplitude of FRET signal was smaller for sCD95L, while for a similar amplitude, the FRET increase was much faster with sCD95L than with IZ-sCD95L. In summary, time-resolved FRET measurements showed that multimerization of receptors by ligand is quick in contrast to FADD recruitment kinetics and that sCD95L has a different receptor crosslinking mechanism than IZ-sCD95L.

## Discussion

In this work we aimed at characterizing how CD95 receptor activation is controlled by the multimericity of ligand and receptor. To this aim, we quantified the receptor and its activity at the single cell level using different fluorescence microscopy approaches. Using FRET acceptor photobleaching we observed no signature of significant oligomerization of CD95. However, using PALM we observed that CD95 is not purely monomeric. To date no structure is reported for CD95 receptor. Assuming a monomer-dimer mixture by similarity to the TNFR structure (*40*), PALM imaging revealed 58% monomeric and 42% dimeric CD95 on the cell surface in the absence of ligand (**Fig. 7A**). Furthermore, on the one hand, we showed that this CD95 dimerization does not influence CD95L apoptosis signaling, as the CD95(R86S) mutant that cannot bind the ligand does not affect CD95-mediated apoptosis when co-expressed with CD95 wild type. On the other hand, when the whole N-terminal part including the CRD1 was removed spontaneous death was observed at lower expression compared to wild-type receptor. We therefore conclude that the multimerization of the unliganded receptor through the PLAD has a protective role, preventing or controlling spontaneous activation of the receptor. Hence we propose that ligand-trimerized receptors are the signaling unit for CD95-induced apoptosis. This conclusion would be in line with the observation that the signaling form of CD95 trimerizes through its transmembrane domain (*38*). This means that partial CD95 dimerization combined with trimerization through ligand interaction may generate larger networks, but they would not influence CD95 induced apoptosis. It has been reported that secondary multimerization of soluble CD95L using antibodies enhances apoptosis compared to soluble CD95L alone (*28*). This hexameric ligand may induce larger signaling units, with a proximity between receptors different from the one induced by the PLAD. These larger signaling units may facilitate FADD recruitment.

**Fig. 7.**
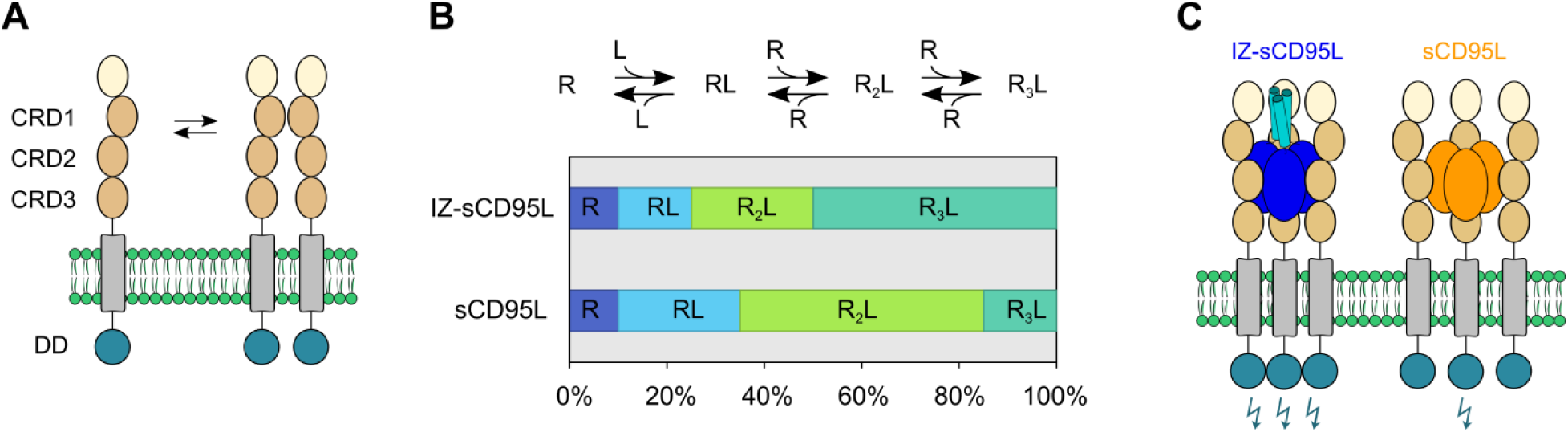
Schematic representation of differential crosslinking of CD95 by sCD95L and IZ-sCD95L. (**A**) At a density of 10 receptors per µm^2^ CD95 exists predominantly as monomer and dimer. (**B**) A possible way to interpret the difference observed by FRET for the two ligands is that their induced receptor trimerization is not complete and that sCD95L generates a lower amount of trimerized receptors than IZ-sCD95L. (**C**) As an alternative to (B) sCD95L and IZ-sCD95L may possess a different structure or rigidity which leads to more or less efficient trimerization.

Quantitative fluorescence microscopy experiments on living cells allowed us to conclude that CD95 receptor activation is a consequence of ligand-induced trimerization of mainly monomeric receptors. Upon addition of sCD95L and IZ-sCD95L, we observed clear signatures for that: the CD95 diffusion coefficient dropped, FRET between receptors appeared and the unbinding of labeled ligand was slower upon washing than upon addition of unlabeled competitor. The soluble form of CD95L is a much weaker inducer of apoptosis than its membrane form (*23*)(*24*). Using dose-response measurements, we observed that sCD95L can induce apoptosis, but even at high concentration sCD95L cannot reach the same activity as its stronger counterpart IZ-sCD95L. From ligand unbinding experiments, sCD95L appeared less stably associated with receptors than IZ-sCD95L. Moreover, FRET measurements between receptors upon ligand binding revealed that sCD95L and IZ-sCD95L trimerize the receptor in a different way, as the correlation between FRET amplitude and kinetics was different. From these observations one can propose that CD95 ligands are able to generate mixes of receptor monomers, dimers and trimers, and that IZ-sCD95L has a stronger capacity to trimerize receptors than sCD95L (**Fig. 7B**). Alternatively, both ligands may oligomerize the receptor with a similar stoichiometry but in a structurally different way, that may result in a different distance or orientation between bound receptors and eventually a different activation (**Fig. 7C**). Both models would be consistent with the different dynamics of diffusion coefficient and FRET signals. The weak apoptotic activity of sCD95L relates to its limited capacity to trimerize the receptor in the active form, but not to its avidity to the receptor. One may hypothesize that this multimerized but weakly apoptotic form of receptor is able to transduce other non-apoptotic signals already observed for sCD95L (*60*)(*61*). Trimerized receptors can only appear from ligand-bound receptors. Thus, from a kinetic perspective, multimerized receptors are not yet present at time zero with a rate of formation that is zero. However we observed on the time scale of minutes that the kinetics of FRET between receptors looked maximal at time zero, showing that receptor multimerization is rapid compared to the first ligand binding. On the contrary, FADD recruitment started with a pronounced delay. Theoretically, a slope of zero at time zero is expected, but a delay of several minutes followed by a more rapid recruitment would not be coherent with a simple, extremely slow interaction model. The observed delay could relate to the time needed for receptors to become active once they are oligomerized, or this may also be explained by the complex biochemistry of FADD recruitment. FADD can self-associate via its DED and the presence of DED has been shown to be important for efficient recruitment to the receptor (*7*)(*8*)(*62*)(*63*)(*64*). Interestingly, the shape of the kinetics of FADD recruitment reminds of the delay time observed for caspase-8 activation (*54*)(*55*)(*65*). One way to interpret this delay consisted in introducing an interaction between caspase-8 on the plasma membrane (*55*). However, our FADD recruitment data let us propose that the delay between receptor clustering and caspase-8 activity rather stems from the DISC assembly itself. We believe that investigation of FADD recruitment and CD95 DD interactions could be future key experiments towards understanding the regulation of CD95 signaling. Fluorescence microscopy now offers many possibilities for the quantification of the stoichiometry of proteins in complexes, in particular with newly developed approaches based on single molecule localization microscopy. These approaches generate a bridge between structural and cellular biology, allowing the *in situ* testing of hypothesis derived from structural studies.

## Materials and Methods

### Plasmids and reagents

Gene knock-out cell lines were generated using CRISPR/Cas9 as in (*66*). For CD95^KO^, the guide RNA was CATCTGGACCCTCCTACCTC, for FADD knock-out, GTTCCTATGCCTCGGGCGCG. CD80, CD86, CTLA-4(Δ23) fusions to mGFP and mCherry were made by replacing mEos2 in the mEos2 fusions described in (*46*) by EGFP-A206K-L221K (*67*) and mCherry (*68*). CD95-mEos2, CD95-mCherry and CD95-mGFP were cloned as the preceding constructs, with GGGGGPVPQWEGFAALLATPVGGAV as linker. For the bicistronic constructs with the 2A peptide, the linker between the two proteins was coding for SGLGSGEGRGSLLTCGDVEENPGPRSRVAT.

CD95 truncations and mutants were generated by PCR, except CD95-R86S that was made by mutagenesis (QuikChange II Site-Directed Mutagenesis Kit, Agilent). The ALPS-Pt2 mutant lacking amino acids 51 to 96 was cloned by amplifying the flanking sequences by PCR and by fusing them with the BsmBI restriction site to prevent scars. CD95-ΔDD corresponds to the first 210 amino acids of human CD95 including the signal peptide.

mCherry-sCD95L and mCherry-IZ-sCD95L were cloned as the mGFP fusions described in (*66*), with the same linkers. In unlabeled IZ-sCD95L, mCherry was replaced by amino acids PS, as described in (*66*). In unlabeled sCD95L, mCherry and the linker LGGGGSG were replaced by amino acids SGR. The caspase-8 activity reporters NES-ELQTDG-mGFP and NES-ELQTDG-EBFP2 were described in (*54*). FADD was fused to mGFP through the linker PRARDPTSGGGGGPVAT and cloned in pIRES-Neo3 (Clontech).

FADD was recognized on western blot by the antibody 2782 (RRID: AB_2100484, Cell Signalling) and HRP-conjugated anti-rabbit secondary antibody (Dianova). CD95 was recognized with the Apo-1-3 antibody (RRID: AB_10541744, Enzo Life Sciences) and an Alexa488 anti-mouse secondary antibody (Dianova).

To calibrate mGFP and mCherry signals, we used mCherry-BID-SNAP-mGFP, designed by fusing mCherry with the linker GGGGSGGGGRVGGGSRG to human BID, followed by the linker GSRAQASNSAVELKLDIT, the SNAP tag, the linker PAGDPPVAT and mGFP.

### Cell lines

HeLa (RRID: CVCL_1922), HT-1080 (RRID: CVCL_0317), LN-18 (RRID: CVCL_0392), the derived knock-out lines and 293T cells were maintained in Dulbecco modified eagle medium containing 10% fetal calf serum, penicillin/streptomycin, 100 μg/ml each (all Thermo Fisher Scientific). HeLa(CD95-mGFP) and HeLa(CD95) are described in (*69*) and (*70*), respectively. Cells were transfected with FuGENE6 (Promega). Stable expression of FADD-mGFP was achieved by selection with G418 (Thermofisher). Methyl-β-cyclodextrin (mβcd) was from Sigma-Aldrich, propidium iodide from Thermofisher. For immunofluorescence, cells were fixed with 4% Pierce™ formaldehyde (16% stock solution, Thermofisher).

### Flow cytometry and fluorescence microscopy

Flow cytometry was performed using the FC500 from Beckman Coulter. Live cell imaging and FRET was done on the SP5 confocal microscope from Leica Microsystems as in (*54*). For FRET measurement by acceptor photobleaching, we measured the mGFP and mCherry intensities before and after mCherry photobleaching. To take into account that not all mCherry molecules are photobleached, we calculated FRET as follows:

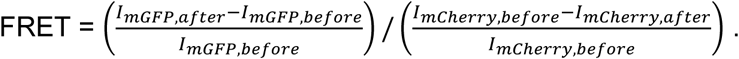

For FRET measurement by sensitized emission, the ratio of fluorescence detected between 600 nm and 670 nm, and between 500 nm and 560 nm was calculated. FRAP and Fluorescence Correlation Spectroscopy (FCS) measurements were performed on the SP2 confocal microscope from Leica Microsystems. FCS was measured using a water immersion objective (HCX PL APO 63 × 1.2 W~CORR), a 594 nm laser line and an FCS-unit (Leica Microsystems, Mannheim, Germany).

### Single-molecule localization microscopy

For quantitative PALM imaging of CD95-mEos2, a custom-built setup was used which is described elsewhere (*46*). Briefly, a 568 nm laser (Sapphire 568 LP, Coherent) and a UV laser (405 nm, Cube 405-50C, Coherent) were focused on the back focal plane of an Olympus IX-71 inverted microscope equipped with a 100× oil immersion objective (PLAPO 100× TIRFM, NA ≥ 1.45, Olympus) and appropriate dichroic mirrors (AHF). A ‘nose piece’ (Olympus) was mounted onto the objective which kept the distance between the sample and objective constant and minimized drift. Fluorescence emission was detected on an EMDDC camera (iXon3 and iXon Ultra, Andor). Recording was started prior to illumination with 568 nm and UV light. Imaging was performed in total internal reflection (TIR) mode with a frame rate of 10 Hz, under continuous 568 nm laser illumination (0.5 kW cm-²) and increasing UV illumination (0-10 Wcm-²) until no more blinking events were observed.

### Quantitative analysis if single-molecule localization microscopy data

Super-resolved images were reconstructed using the rapi*d*STORM software (*71*) and the LAMA package (*72*), following a quantification procedure that was described in detail recently (*73*). In brief, the number of re-activation (“blinking”) events of mEos2 fluorophores in single, super-resolved receptor clusters was determined and plotted in a histogram. The oligomeric state was extracted by approximating the experimental frequency distribution of the number of blinking events by fitting functions derived for monomeric and dimeric receptor clusters (*47*): the monomer function (*p*_0_) is shown in Equation 1,

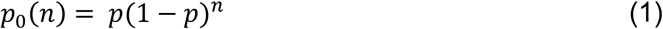

with *n* as the number of re-activation events and *p* as the fraction of mEos2 molecules which do not undergo blinking after photoactivation.

The dimer function (*p*_1_) is shown in Equation 2:

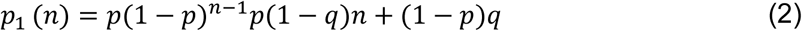

with the additional parameter *q* that corrects for the fraction of mEos2 molecules that were not detected.

Mixed populations of monomer and dimer were analyzed by a linear combination of the monomer and the dimer function (Equation 3),

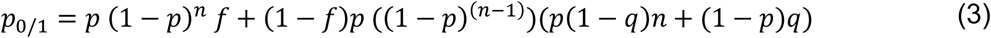

with *f* as the fraction of monomeric proteins within the mixed population (*39*).

## Detailed Attribution of Authorship

C.L. and J. Beaudouin performed experiments, analyzed data and wrote the manuscript. J. Berndt performed experiments and analyzed data. S.A. performed experiments. F.F. and M.H. performed PALM experiments and analyzed data. All authors helped writing the manuscript.

## Acknowledgements

R.E., C.L., J. Berndt and J. Beaudouin thank the Initiative and Networking Fund of the Helmholtz Association within the Helmholtz Alliance on Systems Biology/SBCancer. F.F. and M.H. were supported by the German Science Foundation (DFG, SFB807 and HE/6166-11). We thank Julia Peukes, Sarah Kaspar and Patricia Sauer for technical support.

## Competing financial interests

The authors declare no competing financial interests.

